# *In vitro* evaluation of antifungal activity of *Ageratum conyzoides*, *Basella alba* and *Mitracarpus villosus* on the strain of *Lasiodiplodia theobromae* in the Kisangani Region / D R Congo

**DOI:** 10.1101/2020.12.09.418350

**Authors:** J.T.K. Kwembe, M.K. Asumani, O. Onautshu, P.T. Mpiana, G. Haesaert

**Affiliations:** Department of Chemistry, Faculty of Sciences, University of Kisangani, B.P. 2012 Kisangani, R D Congo; Department of Biotechnological Sciences, Faculty of Sciences, University of Kisangani, B.P. 2012 Kisangani, D.R. of Congo; Department of Chemistry, Faculty of Sciences, University of Kinshasa, B.P. 190 Kinshasa XI, D.R. of Congo; Department of Plants and Cultures, Faculty of Biosciences Engineering, University of Ghent, Ghent 9000, Belgium

**Keywords:** Antifungal activity, *Ageratum conyzoides*, *Lasiodiplodia theobromae*, *Basella alba*, *Mitracarpus villosus*, Kisangani, Activité antifongique, *Ageratum conyzoides*, *Lasiodiplodia theobromae*, *Basella alba*, *Mitracarpus villosus*, Kisangani

## Abstract

Following our previous investigations relating to the in vitro evaluation of antifungal activity, this study focused on the demonstration of the inhibitory power of extracts of fresh and dry leaves of Ageratum conyzoides, Basella alba and Mitracarpus villosus on the strain of Lasiodiplodia theobromae, fungus responsible for brown rot of cocoa pod in the Kisangani region. In six repetitions on the Patato dextrose agar medium, the strain of L. theobromae was inhibited up to 68.1% by the aqueous extract of B. alba; 60.0% by the ethanolic extract of A. conyzoides and 55.2% by the ethanolic extract of M. villosus (all obtained from dry leaves). Only the fresh leaves of the aqueous (56.7%) and ethereal (51.1%) extracts of A. conyzoides showed high inhibition percentages compared to those of the extracts of the fresh leaves of B. alba and M. villosus. The extracts of the dry leaves showed high inhibition percentages followed by those of the fresh leaves and finally those of crude extracts after two days of incubation. Thus, in addition to the expected results, the plants studied are all active. These inhibitory powers could be very high for the secondary metabolites of the respective plants.

## RESUME

Faisant suite à nos investigations antérieures relatives à l’évaluation *in vitro* de l’activité antifongique, cette étude s’est focalisée sur la mise en évidence du pouvoir inhibiteur des extraits des feuilles fraîches et sèches d’*Ageratum conyzoides*, de *Basella alba* et de *Mitracarpus villosus* sur la souche de *Lasiodiplodia theobromae,* champignon responsable de la pourriture brune de cabosse de cacao dans la région de Kisangani. En six répétitions sur le milieu Patato dextrose agar, la souche de *L. theobromae* a été inhibée jusqu'à 68,1% par l’extrait aqueux de *B. alba*; 60,0% par l’extrait éthanolique d’*A. conyzoides* et 55,2% par l’extrait éthanolique de *M. villosus* (tous obtenus des feuilles sèches). Seules les feuilles fraîches des extraits aqueux (56,7%) et éthéré (51,1%) de *A. conyzoides* ont révélé des pourcentages d’inhibition élevés par rapport à ceux des extraits des feuilles fraîches de *B. alba* et *M. villosus*. Les extraits des feuilles sèches ont présenté des pourcentages d’inhibition élevés suivis de ceux des feuilles fraîches et enfin de ceux d’extraits bruts après deux jours d’incubation. Ainsi, subsidiairement aux résultats escomptés, les plantes étudiées sont toutes actives. Ces pouvoirs inhibiteurs pourraient être très élevés pour les métabolites secondaires des plantes respectives.

## 1. INTRODUCTION

Over 40% of the world's population, 52% of them in Africa and Asia, benefit from agriculture, which is considered one of the main sources of income [1, 2]. In this sector, the cultivation of *Cacao theobromae* tends to be predominant compared to that of cotton, sugar cane or tobacco. Africa alone today holds almost 70% of world production [3].

However, this cocoa activity is confronted with the fall in harvest due to fungi, in particular *Lasiodiplodia theobromae*. Reported for the first time on cocoa in Cameroon in 1895 [4, 5] and in South America in Ecuador [6], this pathogenic germ can cause the death of the tree over time [7]. The decline of the cocoa tree orchestrated by *L. theobromae* constitutes a permanent danger not only in India [8], Cameroon and Western Samoa, in the Philippines [9], but also in the Kisangani region of the Democratic Republic of Congo. Among the possible solutions to this scourge is common use and high concentrations of synthetic chemicals, which increases the risk of seeing toxic residues in fruits and/or leaves becoming poisonous over time.

Going through history, it has been found that herbal medicines have contributed enormously to the evolution of modern medicine [10, 11]. About 50% of the therapeutic molecules used today are of natural origin (plants), nurseries of antimicrobial compounds and/or inhibitors of antibiotic resistance mechanisms [12]. Over 80% of the African population uses traditional medicine since it is accessible in terms of cost, proximity and abundance [13]. Indeed, several medicinal plants are used for their biological properties, such as *Ageratum conyzoides*, *Basella alba* and *Mitracarpus villosus* [14].

In addition, several works have highlighted the antimicrobial and/or antifungal potential of extracts from medicinal plants, which have proven to be effective and eco-friendly in the fight against phytopathogenic germs [15, 16, 17]. However, there are almost no studies on the use of plants against *L. theobromae*.

This is how we undertook a series of works on the research of medicinal plants and phytomolecules which can inhibit the growth of the *L. theobromae* strain. This is a saving path which goes hand in hand with the principles of the new regulations which discourage the use of synthetic fungicides [18, 19] and limit the potential risks for human health as well as pollution of the environment [20, 21]

This work aims to evaluate *in vitro* the inhibition power of the four extracts (raw, aqueous, ethanolic and ethereal) from the fresh and dry leaves of *A. conyzoides*, *B. alba* and *M. villosus* on the strain of *L. theobromae*.

## 2. MATERIALS AND METHODS

### 2.1. Study environment

This study was carried out in the Kisangani region, the capital of the Tshopo Province in the Democratic Republic of Congo. The city of Kisangani is located at 0°31 ‘north latitude, from the Equator (57 km), 25°11' east longitude from the Greenwich meridian, and 428 meters above sea level [22, 23].

### 2.2. Plants and treatment

The leaves of *A. conyzoides*, *B. alba* and *M. villosus* were collected after being identified by the Herbarium service of the Faculty of Sciences of the University of Kisangani. These leaves were processed in the laboratory, in three different batches in order to obtain the raw extract from the pressed fresh leaves, the extract from the fresh macerated leaves and the extract from the dried macerated leaves. For this, 10 g of fresh vegetable matter or powder of dry leaves were macerated for 48 hours in 50 ml of distilled water for the aqueous extract, 95% ethanol for the ethanolic extract and diethyl ether for the ethereal extract. The macerated filtrates were concentrated in an oven at 40 ° C, due to 2mL of the concentrate from 10mL of the macerated filtrate.

### 2.3. Obtaining the fungal strain

The *L. theobromae* strain was isolated from cocoa pods naturally affected by brown rot. These pods were harvested directly from the tree in the cocoa plantations of Bengamisa (37 km on the Kisangani-Buta highway) and Yangambi (90 km from the city of Kisangani). After washing the pods, a symptomatic piece was removed, washed with 5% bleach and rinsed in sterile distilled water to finally be seeded on Patato dextrose agar (PDA) at 25 ° C in the dark for five days. Transplanting was carried out on PDA at 25 ± 2 ° C under permanent white light. To prevent the growth of bacteria, 100 μL of Ampicillin (50 mg / mL) and 100 μL of Chloramphenicol (50 mg / mL) were added beforehand to each 100 mL of PDA at 45 ° C before solidification.

### 2.4. Evaluation of antifungal activity

The antifungal activity was evaluated on the basis of percentage inhibition (PI) of mycelial growth or reduction of mycelial growth (RCM) of plant extracts on the strain of *L. theobromae*, with six repetitions. 12mL of PDA were poured into each 90mm diameter Petri dish. A line was drawn in advance on the median of the Petri dish, each studied extract was applied to one half and the mycelial explant 5mm in diameter, was placed on the other half at 2.5mm from the midline [24]. Mycelial growth was measured on either side of the midline (Fungal Ray, FR) every 24 hours until the Petri dish was filled. The negative control was carried out under the same conditions but without extract. Ampicillin and Gentamicin were used as positive controls.

The PI calculation was carried out by the formula

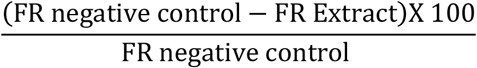

The R 3.4.0 software was used to compare the means of the PI by performing the one-factor variance analysis test (at the 95% threshold). The standard deviation was evaluated in standard deviations represented in error bars on the histograms.

## 3. RESULTS

### 3.1. Inhibition percentage

#### 3.1.1. Extraction solvents

Figure 1 below illustrates the IP of the extraction solvents against *L. theobromae*

**Figure 1:**
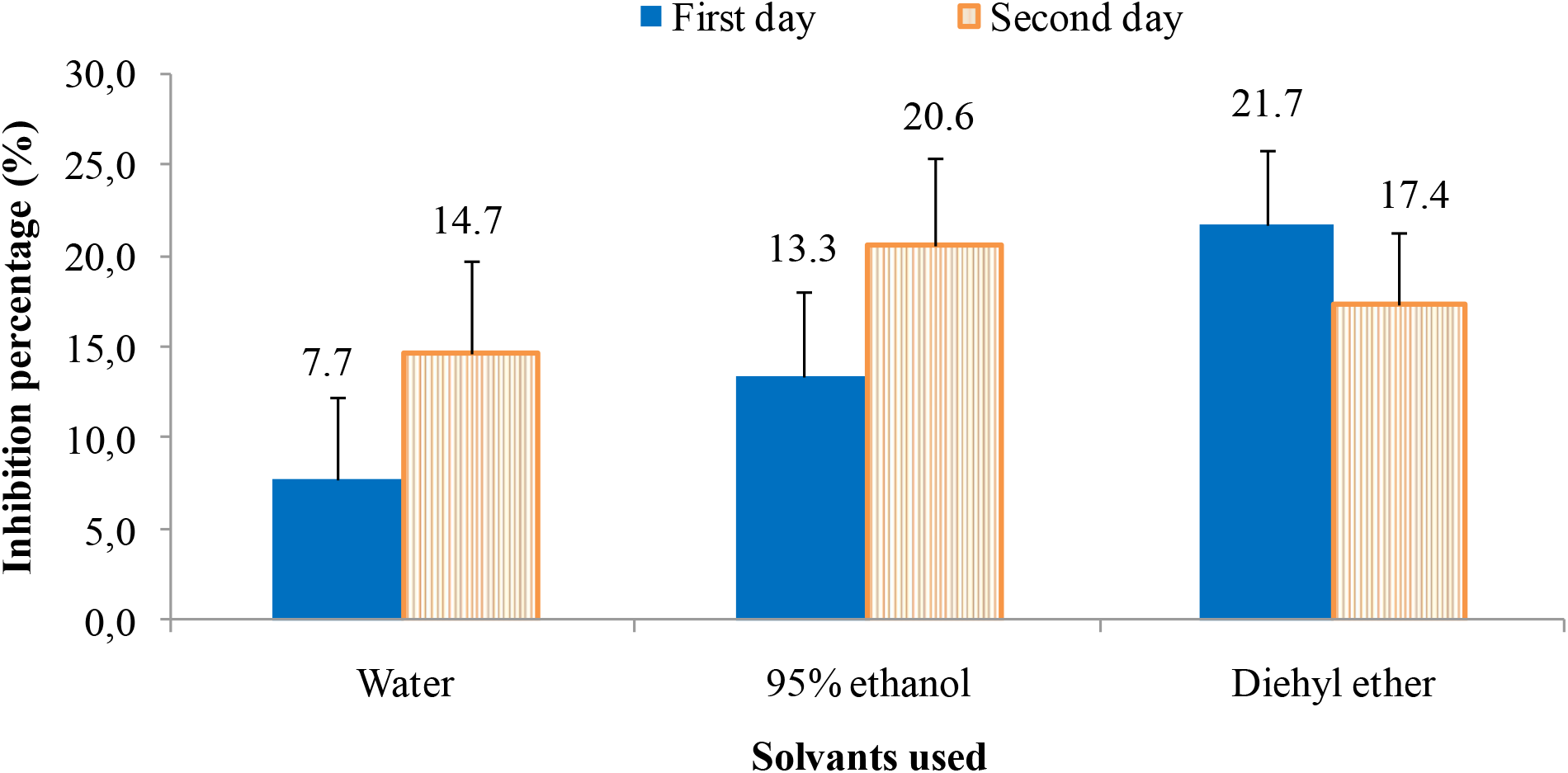
Percentage inhibition of water, 95% ethanol and diethyl ether on the *L. theobromae* strain

After one day of incubation, the diethyl ether achieved a high inhibition rate of 21.7% while the water achieved the lowest PI of 7.7%. However after two days of incubation, the ethanol revealed a high PI at 20.6% while that of the water was low at 14.7%.

#### 3.1.2. Extracts of fresh leaves

Figure 2 below illustrates the antifungal activity of the fresh leaves of the plants tested, after a few days of incubation.

**Figure 2:**
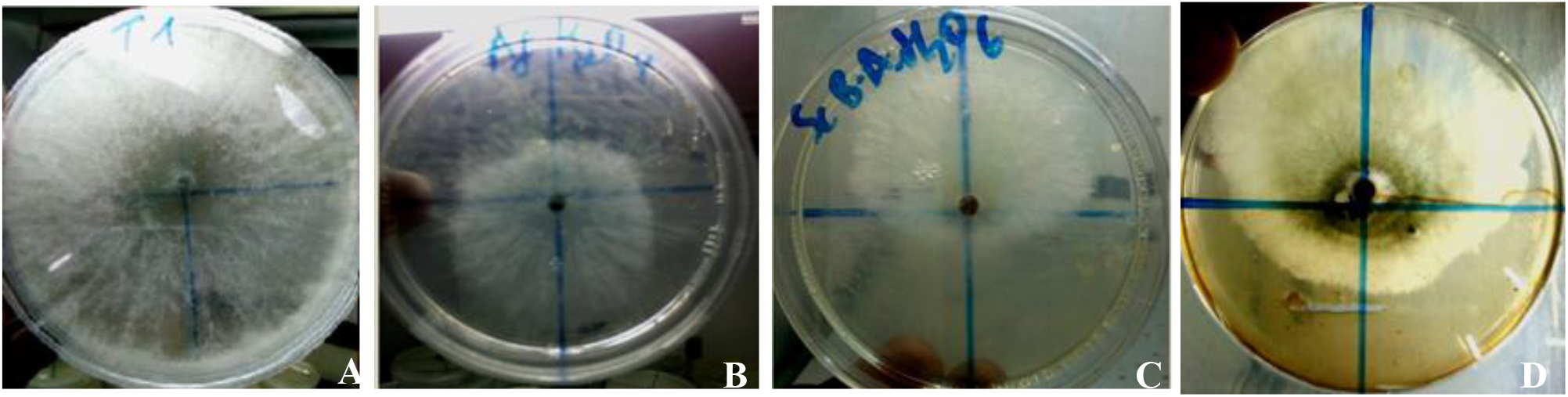
A: Negative control strain of *L. theobromae* (after 2 days of incubation), B: Aqueous extract of fresh leaves of *A. conyzoides* (after 2 days of incubation), C: Aqueous extract of dry leaves of *B. alba* (after 2 days of incubation) and D: Aqueous extract of dry leaves of *M. villosus* (after 4 days of incubation) against the strain of *L. theobromae*

It is obvious to note that the growth of the *L. theobromae* strain was not maximal for *A. conyzoides*, *B. alba* and *M. villosus* whereas it took just 2 days of incubation to achieve maximum growth for the negative control.

##### a. Raw extracts

The fresh leaves of the plants studied revealed PI values as illustrated in figure 3 below.

**Figure 3:**
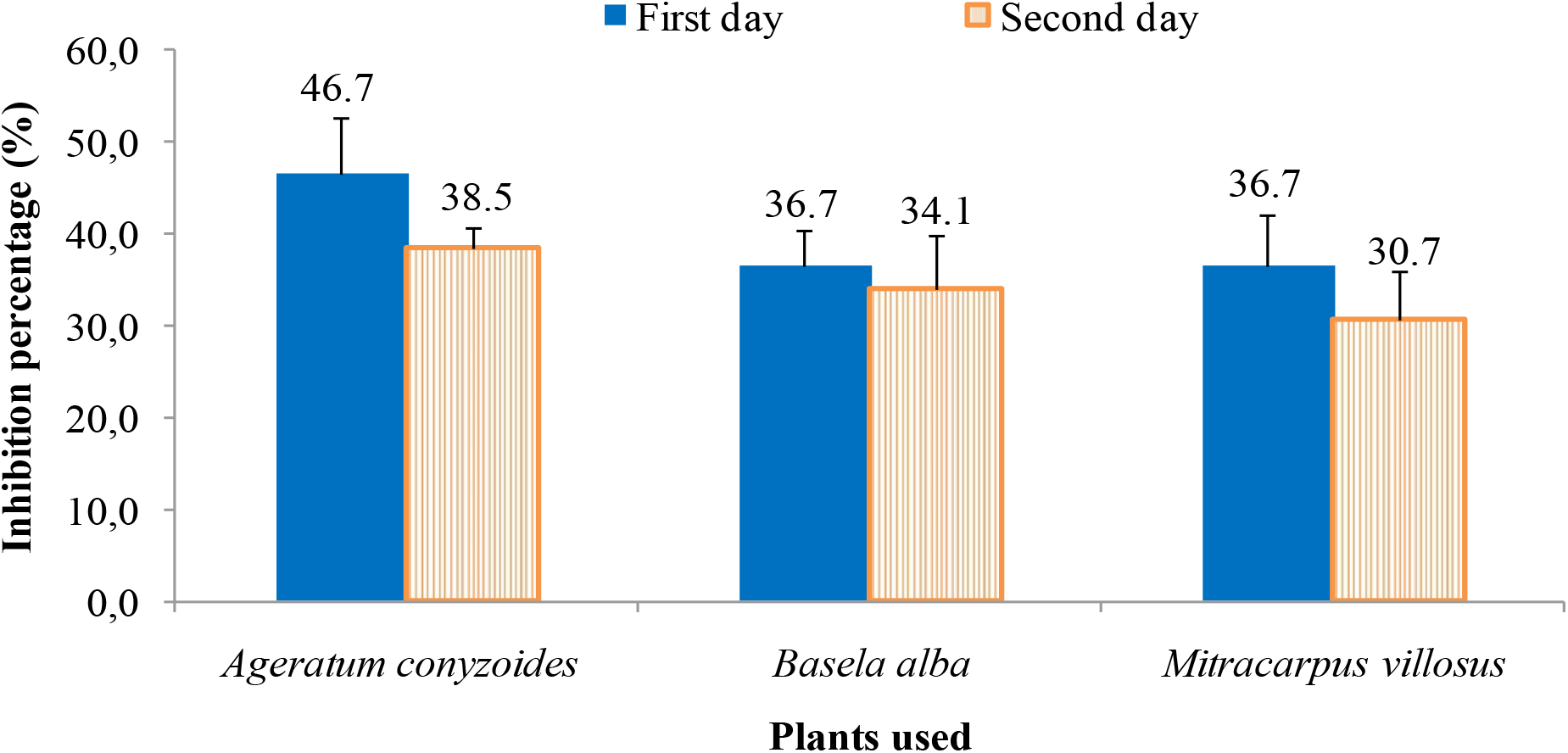
Percentage inhibition of raw extracts of fresh leaves of *A. conyzoides*, *B. alba* and *M. villosus* on the strain of *L. theobromae*

The raw extract of fresh leaves of *A. conyzoides* has high PI during the two days of incubation, 46.8 and 38.5% respectively, whereas that of *M. villosus* had the lowest PI after two days, at 30.7%.

##### b. Macerated extracts

The maceration of the fresh leaves in water, 95% ethanol and diethyl ether revealed the PI on the strain of *L. Theobromae* as illustrated in figure 4 below.

**Figure 4:**
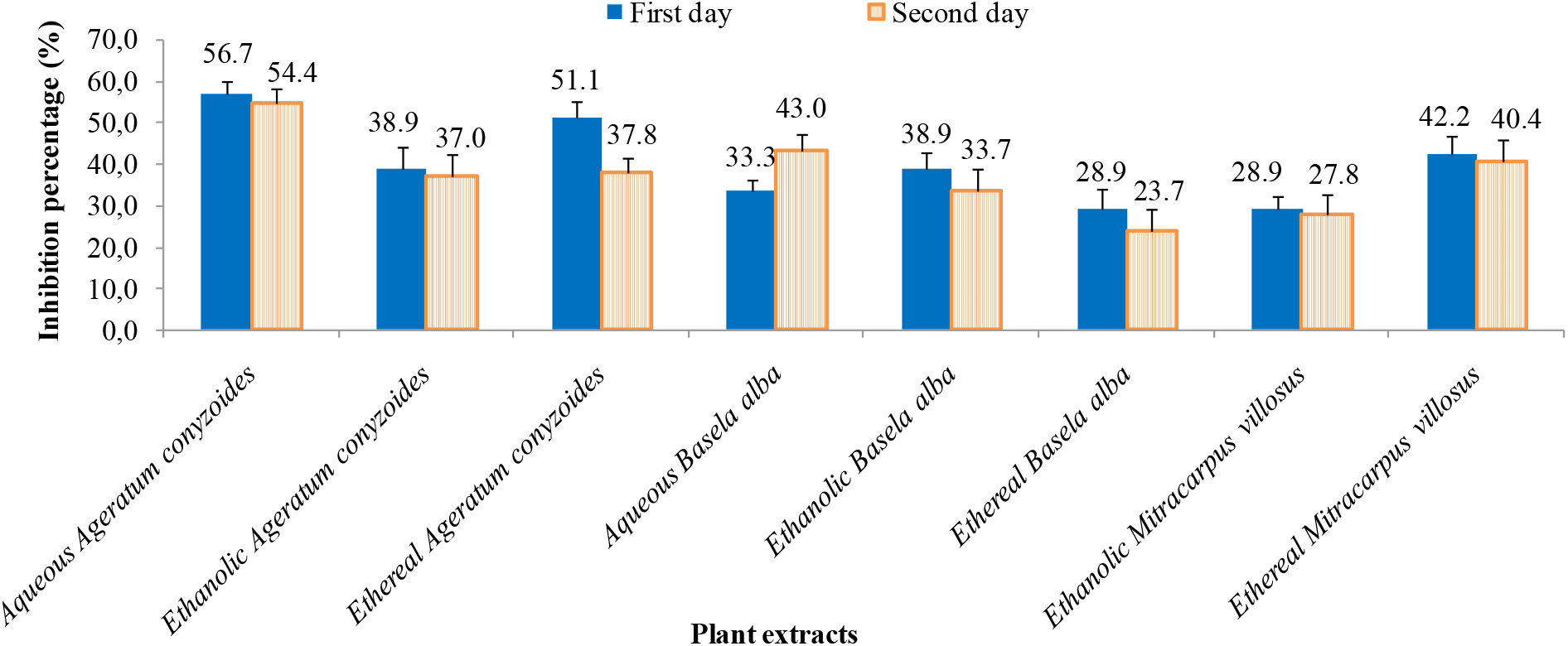
Percentage inhibition of extracts of fresh leaves of *A. conyzoides*, *B. alba* and *M. villosus* on the strain of *L. theobromae*

From this figure, it should be noted that the aqueous extract of *A. conyzoides* has higher PI after one day or 56.7% and two days of incubation or 54.4%; followed by the ethereal extract of the same plant after a day of incubation (51.1%). The ethereal extract of *B. alba* had the lowest PI after two days of observation, 23.7%.

#### 3.1.3. Extracts of dry leaves

The following figure 5 illustrates the PI of the aqueous, 95% ethanolic and ethereal extracts of the dry leaves tested *in vitro* on the strain of *L. theobromae*.

**Figure 5:**
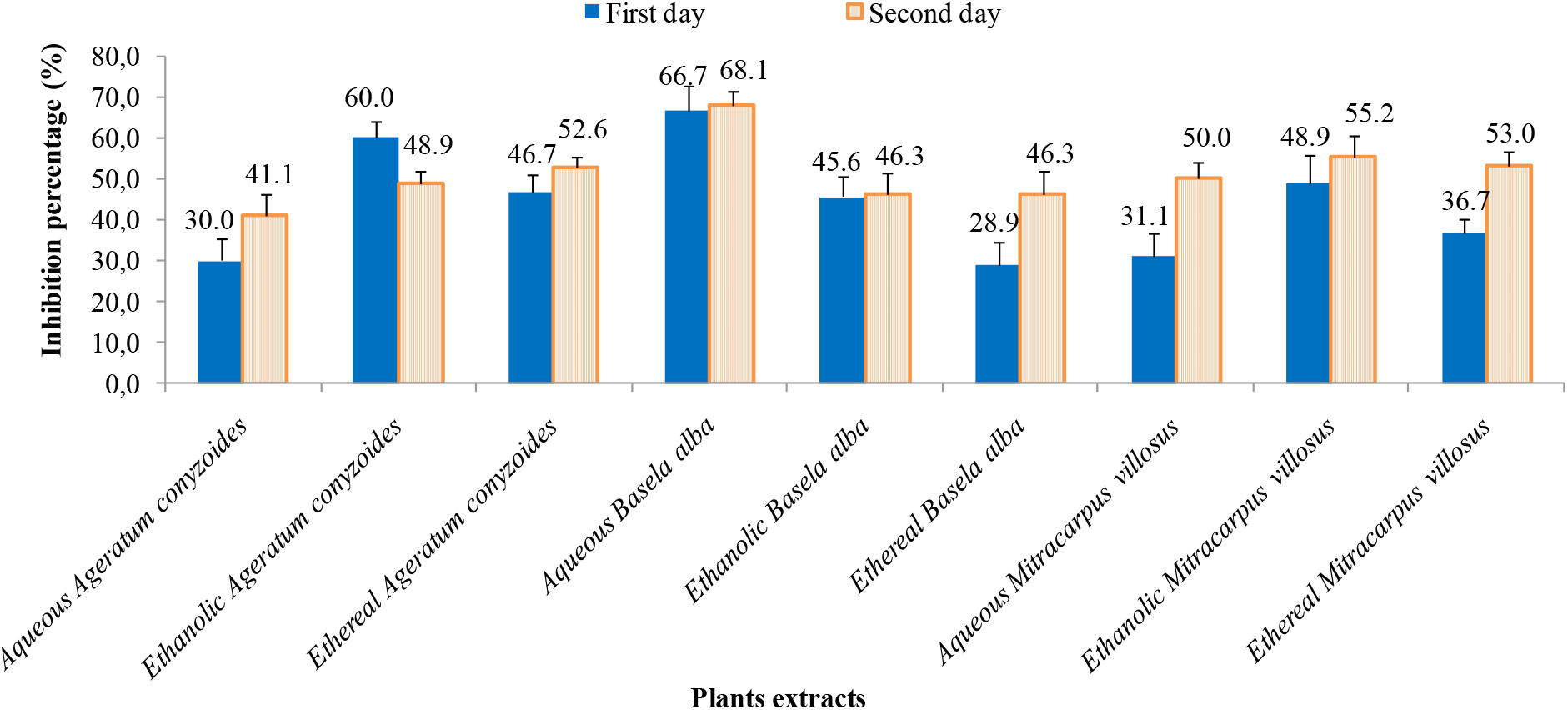
Percentage inhibition of extracts of dry leaves of *A. conyzoides*, *B. alba* and *M. villosus* on the strain of *L. theobromae*

It is obvious to observe from this figure that the aqueous extract of *B. alba* revealed a higher PI after two days of testing 68.1% (66.7% after one day), followed by the 95% ethanolic extract of *A. conyzoides* after one day of incubation, i.e. 60.0%. The ethereal extract of *B. alba* was found to have the lowest PI at 28.9% after one day of incubation.

### 3.2. Maximum growth time

The *L. theobromae* strain is supposed to reach its maximum growth by completely filling the Petri dish. Thus the times necessary to achieve this, called the maximum growth time MGT (in days) for each extract tested are shown in tables 1 and 2 below.

**Table 1:** Maximum growth time (in days) of the *L. theobromae* strain against extraction solvents and antibiotics (positive controls)

| Extraction solvent | Mycelial growth time (day) |
| --- | --- |
| Negative control | 2 |
| Distilled water | 3 |
| Ethanol 95% | 3 |
| Diethyl ether | 3 |
| Ampicillin | 4 |
| Gentamicin | 4 |

**Table 2:** Maximum growth time of the *L. theobromae* strain against the extracts of the plants studied

| Plants used | Mycelial growth time (day) |  |  |  |  |  |  |
| --- | --- | --- | --- | --- | --- | --- | --- |
|  | Raw extract | Aqueous extract |  | Ethanolic extract |  | Ethereal extract |  |
|  | Fresh leaves | Fresh leaves | Dry leaves | Fresh leaves | Fresh leaves | Fresh leaves | Dry leaves |
| *Ageratum conyzoides* | 4 | 4 | 4 | 3 | 4 | 3 | 4 |
| *Basela alba* | 3 | 4 | 6 | 4 | 3 | 3 | 4 |
| *Mitracarpus villosus* | 3 | - | 6 | 3 | 4 | 3 | 4 |

The MGT of the *L. theobromae* strain is only 2 days for the negative control, 3 days for the solvents and 4 days for the antibiotics (Tab 1) like certain plant extracts. On the other hand, with the aqueous extracts of *B. alba* and *M. villosus* (Tab 2), it took 6 days to reach maximum growth.

### 3.3. Water content

Figure 6 below illustrates the water contents of the fresh leaves of the plants studied after their drying.

**Figure 6:**
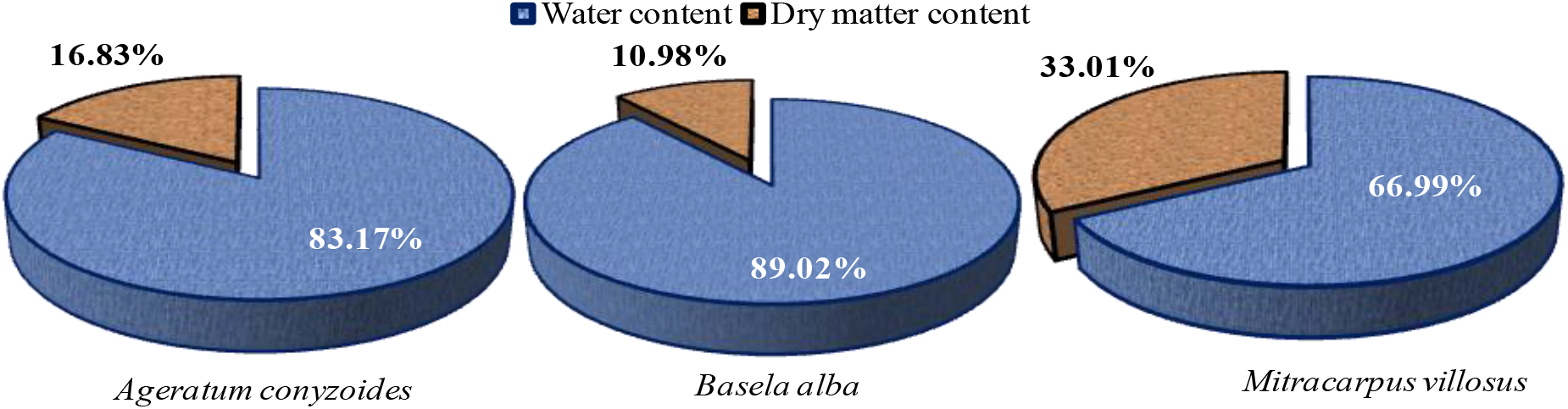
Water content of fresh leaves of *A. conyzoides*, *B. alba* and *M. villosus*

It emerged from this figure that the fresh leaves of *B. alba* are the richest in water, at 89.0% while those of *M. villosus* contain less, at 67.0%.

### 3.4. Total extract content

The extraction yields after complete evaporation of the solvents from the maceration of the dried leaves are illustrated in the following figure 7

**Figure 7:**
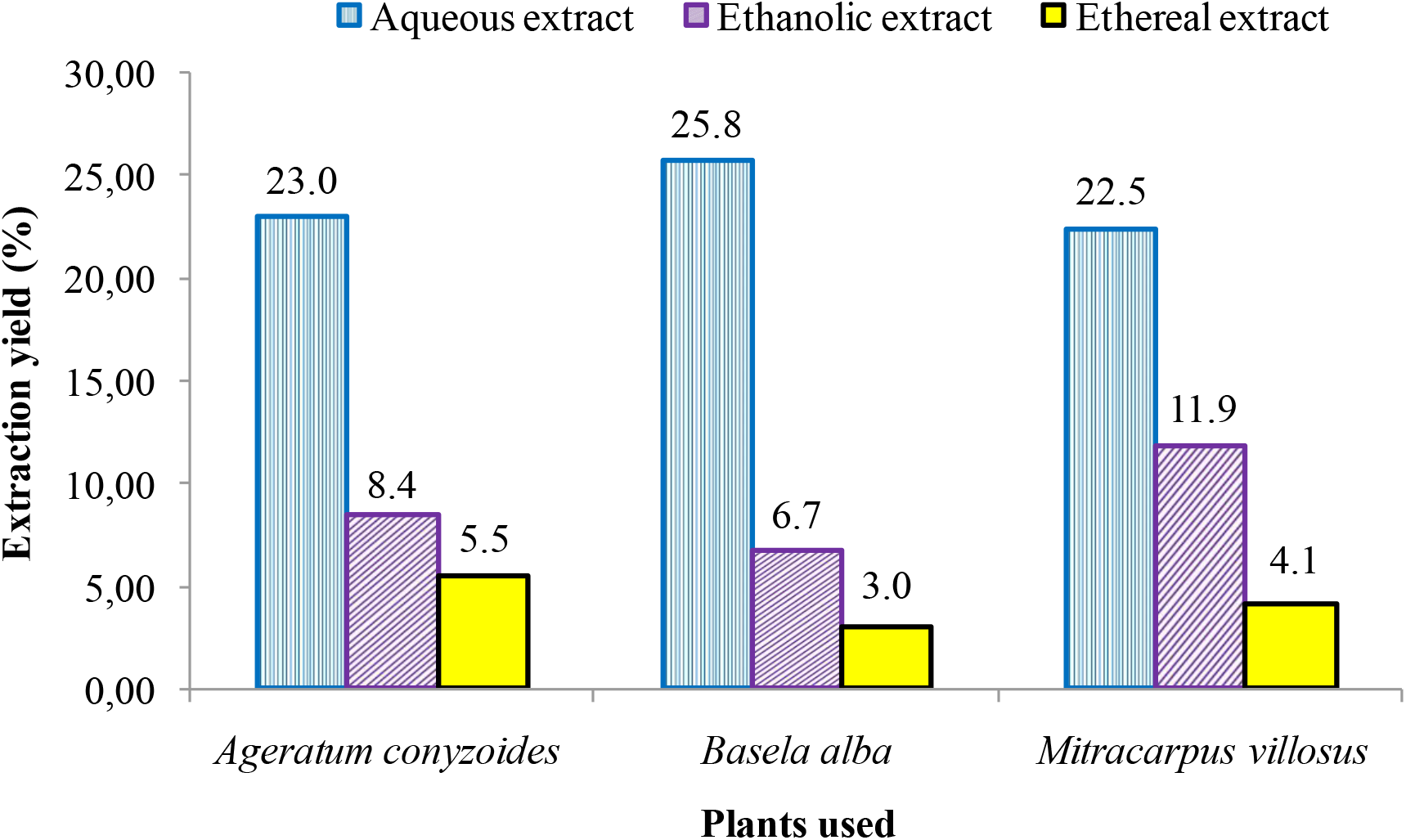
Yield in total extracts

The yield of aqueous extraction of *B. alba* is the highest at 25.7% followed by that of *A. conyzoides* at 23.0%. Furthermore, the ethereal extraction yield of *B. alba* is the lowest, at 2.9%.

## DISCUSSION

As shown in our previous work, the growth of the *L. theobromae* strain was considerably slowed down when confronted to the extracts of the medicinal plants studied (Fig. 2). A prolongation of GMT was found significantly for the tests with extracts, making them pass from 2 days of incubation for the negative control to 6 days in particular for the aqueous extracts of the dry leaves of *B. alba* and *M. villosus*. This same GMT for 6 days was observed for the raw extract of *M. oleifera* [25].

It has been proven that all the solvents used for the extractions (water, 95% ethanol and diethyl ether) have PI showing significant differences in them (p value = 0.00335) but have no antifungal activity (PI less than 25%) according to Bouazza [24]. Furthermore, although ethanol is reputed to be an ideal disinfectant, nonetheless it has been qualified to be ineffective against *L. theobromae*.

The raw plant extracts all have high antifungal activities compared to the negative control, with PI values ranging from 30.7% on the second day of incubation for *M. villosus* to 46.7% on the first day for *A. conyzoides*. Furthermore, the latter has a high inhibitory effect compared to those of *Aloe vera*: 29.6% on the second day and *Newbouldia laevis*: 36.3% on the second day, but has a lower PI than that of *M. oleifera* (74, 1%) after two days of incubation [25]. The raw extracts have significantly different PI (p value = 0.00552 on the first day and p value = 0.0348 on the second day).

In general, the macerated fresh leaves of the plants studied have high PI on the first day of incubation but slightly low on the second day, with the exception of the aqueous extract of *B. alba* which goes from 33.5 to 43.0%. The opposite has been observed for maceration of dry leaves for which the PI generally increase over time, except for the ethanolic extract of *A. conyzoides* which go from 60.0 to 48.9%. This observation would be justified by the presence of very volatile compounds which would constitute the maceration of fresh leaves, in particular essential oils, which mean that over time, these extracts can lose their antifungal power to the advantage of mycelial growth. As for the maceration of dry leaves, this observation would be due to the presence of non-volatile compounds.

Thus it would be preferable to note the results of evaluation of the antifungal activity of volatile compounds (essential oils for example) on the first and second day of incubation, but only on the second day for non-volatile compounds (tannins, alkaloids, saponins, etc.)

The PI of macerated dry leaves are all higher than those of macerated fresh leaves. This would be justified by the fact that in the dry leaves, the extractable matter is in greater quantity than in the fresh leaves following the evaporation of water which the latter contained before drying [25].

It is also necessary to indicate the essential role played by extraction solvents in the selection of the compounds contained in the plants studied. This is the case, for example, of *B. alba*, whose aqueous extract of the dry leaves has a PI of 68.1% on the second day, while the ethanolic and ethereal extracts produced 46.3% inhibition each. There have been very significant differences between different extracts when they are all from the same plant.

Fontem had reported that *M. villosus* (type of extract not reported) had a 64% PI on *Colocasia esculenta* [26]. In the present work, the 95% ethanolic extract of the dry leaves of *M. villosus* has a PI of 55.2% on *L. theobromae* on the second day. This decrease in PI is due to the high virulence of *L. theobromae* compared to *Colocasia esculenta*.

Manal had observed a PI of 28%, for a concentration of 8mg/mL of aqueous extracts of *Lavandula officinalis*, 41% for *Thymus vulgaris*, 43% for *Cymbopogon citratus* against *Botrytis cinerea*, fungus responsible for gray rot of tomatoes in Morocco [27]. All these values are low compared to the PI of aqueous extract of the dry leaves of *B. alba* on the second day (68.1%) and those of the aqueous (54.4% on the second day) and ethereal (51.1% on the first day) extract of fresh leaves of *A. conyzoides*.

Macerated fresh leaves of *A. conyzoides* have higher PI, notably the aqueous extract, 56.8% and 54.4% respectively on the first and second day, followed by ethereal extract on the first day, 51.2%, while the antifungal activity of the raw extract of *A. conyzoids* on the first day was 46.7%. This plant is of fungistatic interest being among the active plants if one takes into account its PI which are between 50 and 75% [24]. Indeed, the latter would contain bioactive molecules because it is cited in several works for its activity on other pathogenic germs. According to Adjou it is used against *Aspergillus flavus*, *Aspergillus parasiticus*, *Aspergillus ochraceus*, *Fusarium oxysporum* [28]. For Djeneb [29] *A. conyzoides* has an antifungal activity against *Candida Albicans*, a zoopathogenic germ while Ndjouondo [30] indicates its role as an antibiotic.

From the standpoint of classification of plants with antifungal activity, in addition to *A. conyzoides*, *B. alba* is also active against *L. theobromae* taking into account the PI threshold observed, ie 68.1% on the second day for the aqueous extract [31, 32]. *M. villosus*, can be considered as an active plant but not at the same level as the first two plants against the fungus studied according the results obtained in present investigation.

The positive controls Ampicillin (42.9%) and Gentamycin (57.8%) all achieved lower PI than that of 95% ethanolic extracts of *A. conyzoides*: 60.0% (on first day) and aqueous extract of *B. alba*: 68.1% on second day (from dry leaves).

As in our previous investigations [25], the yields of aqueous extracts are far higher than those of ethanolic and ethereal extracts. Indeed, on the basis of the "like disolves like" principle, water, which is the universal solvent, generally dissolves a greater number of substances than any other solvent, in particular polar substances [33].

On the one hand, the yield of aqueous extract (23.0%) of *A. conyzoides* is higher compared to that obtained by Adjou, ie 15.01% [28], but for the same plant, its yield of ethanolic extract (23.58%) is higher than that found in this study (8.4%) on the other hand.

According to the results obtained by Kambou [34], *Mitracarpus scaber* has a yield of 4% for the ethanolic extract, a low yield compared to that of *M. villosus* in the present work (11.9%), but that of aqueous extract (5.3%) of *M. scaber* is higher than that of *M. villosus* (4.1%). These differences would depend on the species of the plants studied.

*Mitracarpus scaber* has a yield of ethanolic extract (4%) is lower than and aqueous extract (5.3%) obtained by Kambou [34]. All these values are much lower than those of *M. vullos* is whose ethanolic extract yield is 11.10% and that of aqueous extract 20.80%. These differences would be due to the chemical composition of the species of the plant studied.

The water contents of *B. alba* (89.02%) and *A. conyzoides* (87.34%) are all higher than those of *M. oleifera* (74.2%) [22] and even that of *M. oleifera* found by Broin, 75% [35], of *N. laevis* (75.3%) but are lower than that of *A. vera* (93.3%) found previously [22]

Tsopmbeng obtained a low yield of aqueous extract (21.20%) of *Cupressus lusitanica* [36] compared to our results of aqueous extract of *B. alba*, (23.16%). It should be emphasized that the aqueous extraction yields of *A. vera* (15.4%) and *M. oleifera* (17.0%) and *N. laevis* (8.1%) [22] are all low compared to those of the plants studied in this work.

From the results obtained, very significant differences emerged between the PI of ethereal (p value = 0.00218), ethanolic (p value = 0.00987) and aqueous (p value = 5.95.e-08) dry leaves. It is therefore important, in our next investigations to make a bio-guided fractionation to identify and extract the phytochemical groups responsible for the antifungal activity of these plants.

## CONCLUSION

The aim of this work was to evaluate, in vitro, the inhibitory power of extracts of fresh and dry leaves of *Ageratum conyzoides*, *Basella alba* and *Mitracarpus villosus* on the strain of *Lasiodiplodia theobromae* isolated from brown cocoa pod. It appears that all the extracts of the plants studied are active and have a fungistatic character according to the method used.

The high percentages of inhibition were observed for the extracts of dry leaves including that of the aqueous extract of *B. alba* (68.1% after two days of incubation), of the ethanolic extract of *A. conyzoides* (60, 0% after one day of incubation) and of the ethanolic extract of *M. villosus* (55.2% after two days of incubation). The extracts of the fresh leaves revealed average percentages of inhibition, in particular that of the aqueous extract of *A. conyzoides* (56.7 and 54.4% respectively after one and two days of incubation) and that of the ethereal extract from the same plant (51.1% after one day of incubation). A more dominant threshold of percentage inhibition could be observed for the secondary metabolites contained in these medicinal plants. Thus, our subsequent work will focus on the identification and isolation of the main phytochemical groups of these plants in order to assess their respective antifungal activities on the strain of *L. theobromae*.

## ACKNOWLEDGEMENT

Professor Geert Baert of the University of Ghent/Belgium and North Coordinator of VLAAMSE INTERUNIVERSITAIRE RAAD (VLIR-UOS)/University of Kisangani: for assistance in the acquisition of some laboratory equipment and especially during an internship at the University of Ghent.

NKFUTELA EWALA Michel for his scientific and deontological contribution.

